# Reactive Oxygen Species-induced Protein Carbonylation Promotes Deterioration of Physiological Activity of Wheat Seeds

**DOI:** 10.1101/2022.01.24.477486

**Authors:** Bang-Bang Li, Shuai-Bing Zhang, Yang-Yong Lv, Shan Wei, Yuan-Sen Hu

**Author notes:** Corresponding Author. College of Biological Engineering, Henan University of Technology, Lianhua Street, Zhengzhou, Henan, 450001, PR China. E-mail address (Y.-S. Hu).

## Abstract

During the seed aging process, reactive oxygen species (ROS) can induce the carbonylation of proteins, which changes their functional properties and affects seed vigor. However, the impact and regulatory mechanisms of protein carbonylation on wheat seed vigor are still unclear. In this study, we investigated the changes in wheat seed vigor, carbonyl protein content, ROS content and embryo cell structure during an artificial aging process, and we analyzed the correlation between protein carbonylation and seed vigor. During the artificial wheat-seed aging process, the activity levels of antioxidant enzymes and the contents of non-enzyme antioxidants decreased, leading to the accumulation of ROS and an increase in the carbonyl protein content, which ultimately led to a decrease in seed vigor, and there was a significant negative correlation between seed vigor and carbonyl protein content. Moreover, transmission electron microscopy showed that the contents of protein bodies in the embryo cells decreased remarkably. We postulate that during the wheat seed aging process, an imbalance in ROS production and elimination in embryo cells leads to the carbonylation of proteins, which plays a negative role in wheat seed vigor.

## Introduction

Wheat (*Triticum aestivum* L.), an important food grain, plays an irreplaceable role in making noodles, steamed buns, fodder, alcoholic beverages, biofuel and a variety of other products [1–2]. Consequently, the quality of seeds plays an important role in agricultural development and food quality. Post-harvest wheat grain is commonly stored in warehouses for over 1 year and even longer in China [3]. However, as the storage time increases, or under adverse conditions, the quality of the wheat grain, including vitality and edible quality, inevitably deteriorates. The quality is affected by a variety of environmental factors, especially temperature and humidity. This is often referred to as ageing [4]. Ageing leads to a decrease in the quality of wheat grain, which has an adverse impact on the agricultural production and wheat grain-containing food processing [5]. Therefore, the deterioration of wheat quality during storage has always been a focus of research.

The decrease in wheat seed vigor caused by temperature and humidity during storage is a main result of grain quality deterioration. Postharvest wheat grain is a living organism with a lower metabolic level, and reactive oxygen species (ROS) accumulate in cells under adverse storage conditions, which is the inherent factor that reduces seed vigor [6]. Wheat seeds contain a variety of ROS, including superoxide anions (O_2_^−^), hydrogen peroxide (H_2_O_2_), hydroxyl radical, singlet oxygen and hydroxide ions [7–8]. ROS are extremely active molecules that can oxidize proteins, lipids, DNA and other biological macromolecules in cells, ultimately causing cell death [9–12]. In addition, there are also ROS-scavenging systems in plant cells, including antioxidant enzymes, such as superoxide dismutase (SOD) and peroxidase (POD), and non-enzymatic antioxidants, such as ascorbic acid and glutathione [13–19]. Therefore, there is usually a dynamic balance between the production and removal of ROS. However, with a prolonged storage time and the stress of environmental, this balance is disturbed, and the seed cells are subjected to the oxidative attack of ROS, which eventually leads to seed inactivation [20–21]. The loss of seed viability has been linked to damage caused by ROS [22–23].

Proteins play important physiological roles in cells and their oxidative modifications can seriously affect seed vigor. Protein carbonylation is an irreversible protein oxidation reaction, caused by ROS attacks on proteins, and it is a marker of protein oxidation. ROS can directly oxidize arginine, lysine, threonine, proline and other residues of protein side chains to form carbonyl groups under the catalysis of the transitional metal ion system [24–25]. In addition, ROS can also extract hydrogen atoms from the α-carbon of a protein chain, causing the oxidative hydrolysis of the carbon atoms and the formation of a carbonyl group [26–27]. Moreover, ROS can catalyze lipid peroxidation and non-enzymatic glycosylation to produce some active carbonyl compounds, which are cross-linked to protein side chains through the carbonyl ammonia reaction to form carbonyl groups, and which react further to form carbonyl proteins [28]. The carbonyl proteins cannot be repaired and can only be degraded in a 20S proteasome-dependent manner. However, when the carbonyl protein cannot be eliminated in a timely fashion, it accumulates in the cell, resulting in cell apoptosis [29–31]. Carbonylation affects the synthesis and conformational processes of proteins associated with responses to stresses caused by ageing, and carbonylation is associated with a decrease in the germination capacity [32–33] (Nguyen et al., 2015; Yin et al., 2017).

In this study, ROS production and its impact on protein carbonylation in two wheat varieties with different hardness levels were analyze during an artificial wheat grain aging process. Additionally, the relationship between protein carbonylation and decreased seed vigor was explained through observations of embryonic cellular structures. The data has provided new insights into understanding the deterioration of wheat grain.

## Materials and methods

### Wheat grain and artificial ageing treatment

Wheat grain from of two cultivars, Zhengmai-9023 (ZM 9023) and Yangmai-15 (YM 15), were obtained from the Qiule Seed Company (Zhengzhou, China). The average seed moisture content was adjusted to 11%. Each 200 g sample of seeds was placed in a polyethylene net, stored in a glass drier with a saturated sodium bromide solution at the bottom, and placed in the drier of an oven at 50 °C to ensure that the wheat seeds were at a constant 50 ± 0.2 °C and 50% ± 2% humidity. Samples were withdrawn each week. The wheat embryos were removed with a clean scalpel at 4 °C and sufficiently ground in liquid nitrogen before use.

### Germination potential and germination rate

The germination potentials and germination rates were determined following Chinese National Standard protocol GB/T 5520-2011. A total of 100 wheat seeds were placed in a petri dish (d = 12 cm) with two layers of filter paper (presoaked in distilled water) and placed at 20 °C in an incubator. The germination potentials and germination rates were calculated after 3 and 7 days of incubation, respectively. The wheat seeds were not considered to have germinated if the wheat root was less than half the length of the seed. All the observations were made in triplicate (similarly hereinafter). The germination potentials and germination rates were determined every week.

### Carbonyl protein content

The quantification of carbonyl was carried out in accordance with the method of Boucelha et al [34], with some modifications. Briefly, 0.02 ± 0.001 g wheat embryo was weighed, put in a 1.5-mL centrifuge tube and homogenized with 1.0 mL HEPES extraction buffer (10 mM HEPES-NaOH buffer, 0.1% protease inhibitor cocktail and 0.07% β-mercaptoethanol, pH 7.5) in an ice bath. Samples were centrifuge at 8,000 g for 10 min at 4 °C. Then, two 1.5-mL centrifuge tubes containing 0.2 mL of supernatant were prepared, and 0.4 mL 2-M HCl and 0.4 mL 2-N dinitrophenylhydrazone were added to the control and the experimental groups, respectively. Tubes were incubated for 1 h at 37 °C in the dark. Then, 0.5 mL 40% trichloroacetic acid was added, incubated for 5 min at 4 °C, and centrifuged at 8,000 g for 15 min at 4 °C. The supernatant was discard, and the pellet washed with 1 mL ethanol/ethyl acetate solution (1/1, v/v). Samples were then vortexed and centrifuge again at 8,000 g for 10 min at 4 °C. The sediment was dissolved in 1.0 mL guanidine hydrochloride, tubes were incubated for 15 min at 37 °C, and then centrifuged at 8,000 g for 15 min at 4 °C. In total, 1 mL of supernatant was used to measure carbonyl at an absorbance of 370 nm, and compared with a blank control. The carbonyl protein content was expressed in μmoL/g.

### The O_2_^−^ and H_2_O_2_ contents

The quantification of O_2_^−^ was carried out in accordance with the method of previously reported methods [35], with some modifications. Briefly, 0.02 ± 0.001 g wheat embryo was weighed with 1 mL of 65 mM phosphate buffer (pH 7.8), put in a 2-mL centrifuge tube and ground carefully in an ice bath. They were then centrifuged at 8,000 g for 20 min at 4 °C. Afterwards, 0.2 mL of supernatant was removed, placed in a 2-mL centrifuge tube with 0.3 mL of phosphate buffer and 0.4 mL of 10 mM hydroxylamine hydrochloride and incubated for 20 min at 37 °C. Then, 0.3 mL of 58-mM sulfanilamide and 0.3 mL of 7-mM α-naphthylamine were added to the sample and incubated for 20 min at 37 °C. Afterwards, 0.5 mL of chloroform was added to the sample, mixed and centrifuged at 6,000 g for 5 min at 25 °C. In total, 1 mL of supernatant was absorbed and measured at 530 nm, and a standard curve was established using a sodium nitrite standard solution. The standard curve was used to calculate the O_2_^−^ content (μmoL/g).

For H_2_O_2_ determinations, 0.05 ± 0.001 g wheat embryo was weighed, and the H_2_O_2_ contents were determined using an H_2_O_2_ assay kit in accordance with the manufacturer’s protocol (Solarbio, Beijing, China).

### SOD, catalase (CAT) and POD activity levels

To determine the SOD activity, 0.02 ± 0.001 g wheat embryo was weighed with 1 mL of ice-cold phosphate buffer (pH 7.8), put in a 1.5-mL centrifuge tube and ground carefully in an ice bath. Samples were centrifuge at 8,000 g for 20 min at 4 °C. Then, 90 μL of supernatant was mixed with 240 μL of 50-mM phosphate buffer (pH 7.8), 6 μL of 13-mM methionine, 180 μL of 75-μM nitroblue tetrazolium, 480 μL of deionized water and 30 μL of 0.1-mM EDTA, and then, the reaction was maintained for 30 min at 37 °C. In the control, methionine was not added. Afterwards, 1 mL of the reaction mixture was used to measure the SOD activity at an absorbance of 370 nm, and compared with a blank control. The activity was expressed as units per milligram (U/mg) of wheat embryo.

The CAT activity was assayed as described by Wu et al [36], with some modifications. Samples were processed as above. In total, 35 μL of supernatant was added to 1 mL of 50-mM phosphate buffer (pH 7.8), and the reaction was initiated by the addition of 5 μL of 30% H_2_O_2_. The decrease in the absorbance at 240 nm was measured for 1 min. The catalytic degradation of 1 μmoL H_2_O_2_ per minute per gram of wheat embryo in the reaction system was defined as one unit of enzyme activity.

The quantification of POD activity was assayed as described by Madhava Rao [37], with some modifications. Samples were processed as above. The reaction mixture contained 15 μL of supernatant, 270 μL of deionized water, 520 μL of 125-mM phosphate buffer, 130 μL of 50-mM pyrogallol and 135 μL of 50-mM H_2_O_2_. The increase in the absorbance at 470 nm was measured from 30 s to 90 s. The increase in the absorbance at 470 nm by 0.01 per minute per gram of wheat embryo in the reaction system was defined as one unit of enzyme activity.

### Glutathione content

In total, 0.1 ± 0.001 g wheat embryo was weighed, and the oxidized glutathione (GSSG) content was determined using a GSSG assay kit in accordance with the manufacturer’s protocol (Solarbio).

To determine the reduced glutathione (GSH) content, 0.1 ± 0.001 g wheat embryo was weighed with 1 mL 5% monophosphoric acid and thoroughly ground in an ice bath. The samples were centrifuged at 8,000 g for 10 min at 4 °C. Then, 0.1 mL of supernatant, 0.7 mL of 0.1 M phosphate buffer and 0.2 mL of 4 mM dinitrophenylhydrazone were added to a 1.5-mL centrifuge tube. After mixing, the absorbance value was measured at 412 nm. Deionized water was used instead of the supernatant as the control, and a GSH standard solution with a certain concentration gradient was used to establish the standard curve. The standard curve was used to calculate the GSH content (μmoL/g) [38].

### The ultrastructures of wheat embryo cells

The wheat embryos were artificially aged for 0 and 42 days and then placed in 4% glutaraldehyde solution for pre-fixation for 4 h. They were then rinsed four times, for 15 min each, with phosphate buffer saline at pH 7.2. Afterwards, they were put into a 1% osmic acid solution for a post-fixation of 90 min. They were rinsed with PBS buffer three times, for 15 min each. The samples were successively placed for 15 min into an acetone solution with a concentration gradient of 30%, 50%, 70%, 90% and 100%, for gradual dehydration. Then, they were successively soaked for 120 min in 1:2, 1:1, 2:1 and 1:0 epoxy resin:acetone mixtures. The soaked samples were placed in an air drying oven at 37, 45 and 60 °C and heated for 12 h. Finally, the samples were sliced with an LKB-NOVA ultra-thin slicer to an 80-nm thickness. After staining for 25 min with uranium acetate and 20 min with lemon lead, images were taken under a transmission electron microscope (TEM) [39].

### Statistical analysis

For the germination potentials, germination rates, carbonyl protein, O_2_^−^, H_2_O_2_, GSH and GSSG contents, and the SOD, CAT and POD activity levels, three replicate experiments were conducted. The data were evaluated by applying an analysis of variance and a correlation analysis using SPSS 20.0, and the figures were prepared using GraphPad Prism 6.0.

## Results

### Deterioration of seed vitality

In the early stage of ageing, wheat seeds of both cultivars maintained high germination rates and germination potentials, with fluctuations. However, the germination potentials and germination rates of wheat seeds significant decreased after the artificial aging treatment (Fig. 1), and they began to show downward trends and declined sharply at 14 and 21 days, respectively, of artificial aging. In addition, the germination potentials of the two wheat seed cultivars declined to approximately 0% and 10%, respectively, after approximately 42 days. The germination rates of two wheat seed cultivars decreased to approximately 10% after the artificial aging treatment.

**Fig.1.**
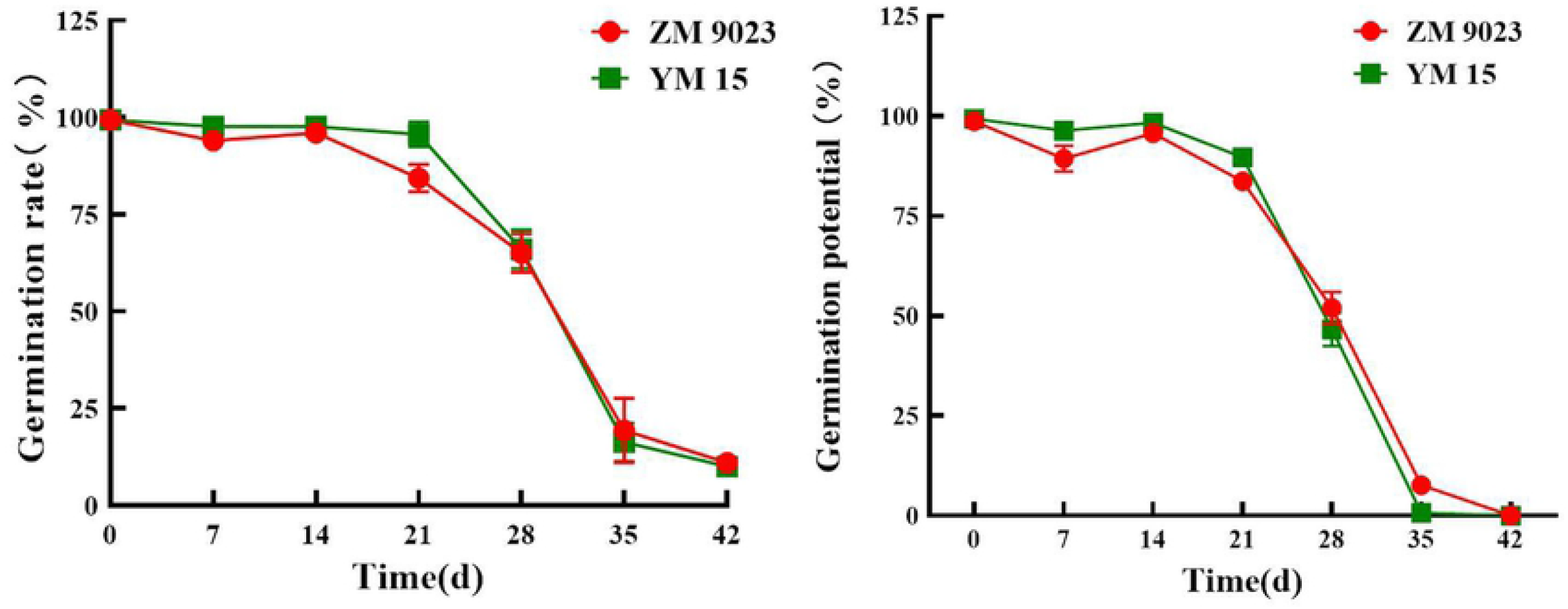
Effects of the artificial aging treatment on wheat seed vigor. Influence of artificial aging treatment on the wheat seed (A) germination potential and (B) germination rate.

### Changes in the carbonyl protein content

The carbonyl protein contents of wheat seeds during storage increased progressively, with some fluctuations, as shown in Fig. 2. The carbonyl protein contents in seeds of YM 15 and ZM 9023 increased by 2.5 and 1.9 times, respectively. The carbonyl protein content of ZM 9023 increased from 0.3077 μmoL/g to 0.5892 μmoL/g after 42 days, and the carbonyl protein content of YM 15 increased from 0.2092 μmoL/g to 0.5228 μmoL/g after 42 days during artificial aging.

**Fig.2.**
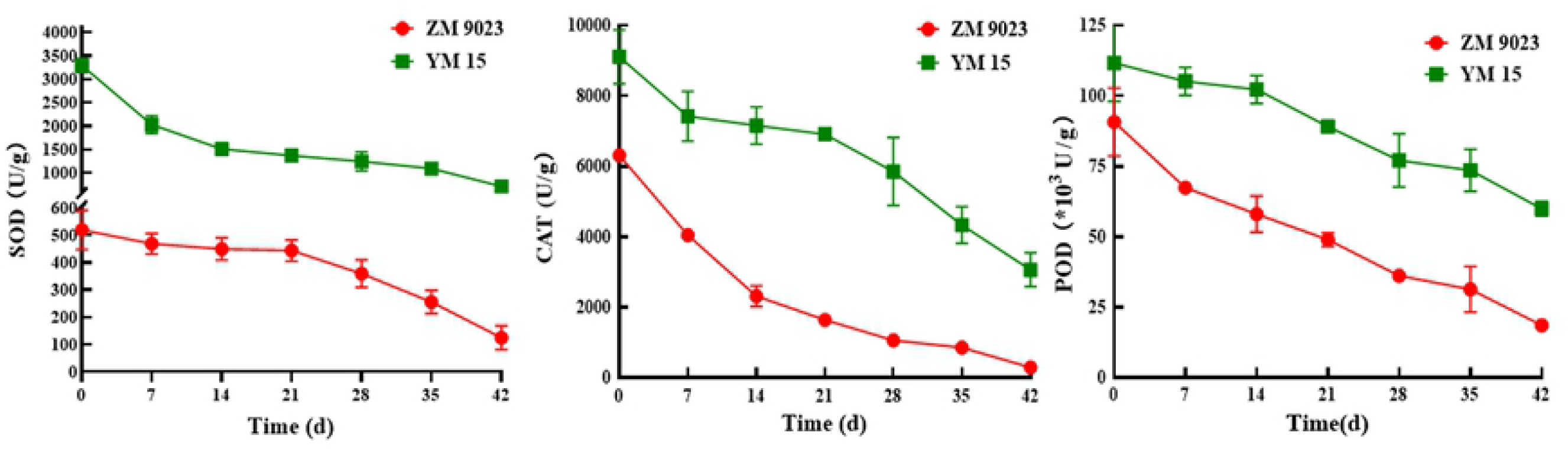
Changes in the carbonyl protein contents of the wheat cultivars ZM 9023 and YM 15 during artificial aging.

### Changes in the ROS content

Both O_2_^−^ and H_2_O_2_ are common ROS in wheat seed cells. Their changes during storage are shown in Fig. 3. The O_2_^−^ contents in both cultivars did not significantly change during artificially ageing. In contrast, the H_2_O_2_ contents of ZM 9023 and YM 15 increased significantly, from 2.70 μmoL/g to 5.53 μmoL/g for the former and from 6.73 μmoL/g to 10.96 μmoL/g for the latter during artificial ageing.

**Fig.3.**
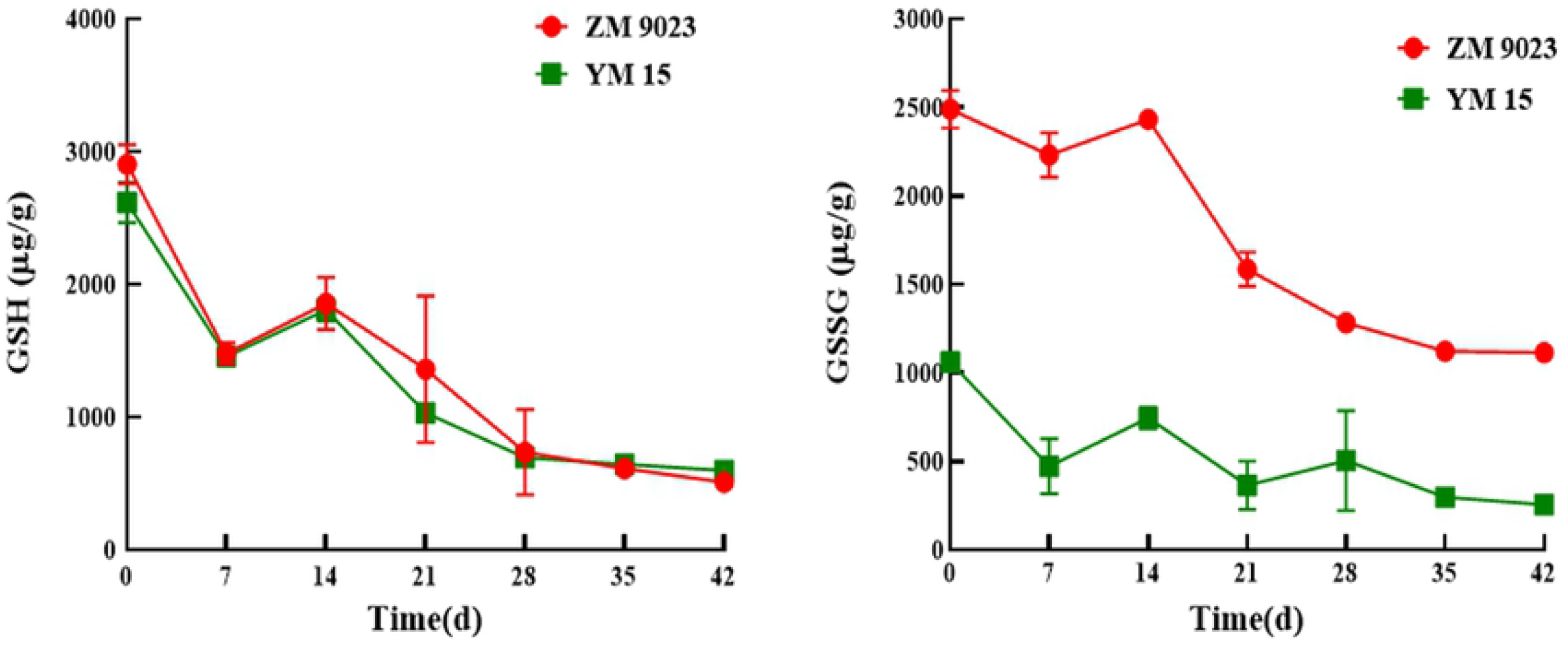
Changes in the superoxide anion (O_2_^−^) and hydrogen peroxide (H_2_O_2_) contents of the wheat cultivars ZM 9023 and YM 15 during artificial aging.

### Changes in antioxidant enzyme activity levels

In general, the activities of antioxidant enzymes in wheat seeds decreased significantly during artificial aging (Fig. 4). The SOD activity levels in ZM 9023 and YM 15 declined 76.55% and 78.37% under the artificial aging treatment, respectively. Additionally, the CAT activity levels of both ZM 9023 and YM 15 decreased significantly, by 21.99 and 2.97 times after the artificial aging treatment, respectively. Furthermore, the POD activity level of ZM 9023 declined from 9.03 × 10^3^ U/g to 1.85 × 10^3^ U/g during artificial aging, and it decreased significantly after 7 days of storage. The POD activity level of YM 15 declined from 11.17 × 10^3^ U/g to 5.99 × 10^3^ U/g during artificial aging, and it decreased obviously after 21 days of storage The POD activity level of YM 15 declined from 11.17 × 10^3^ U/g to 5.99 × 10^3^ U/g during artificial aging, and it decreased obviously after 21 days of storage.

**Fig.4.**
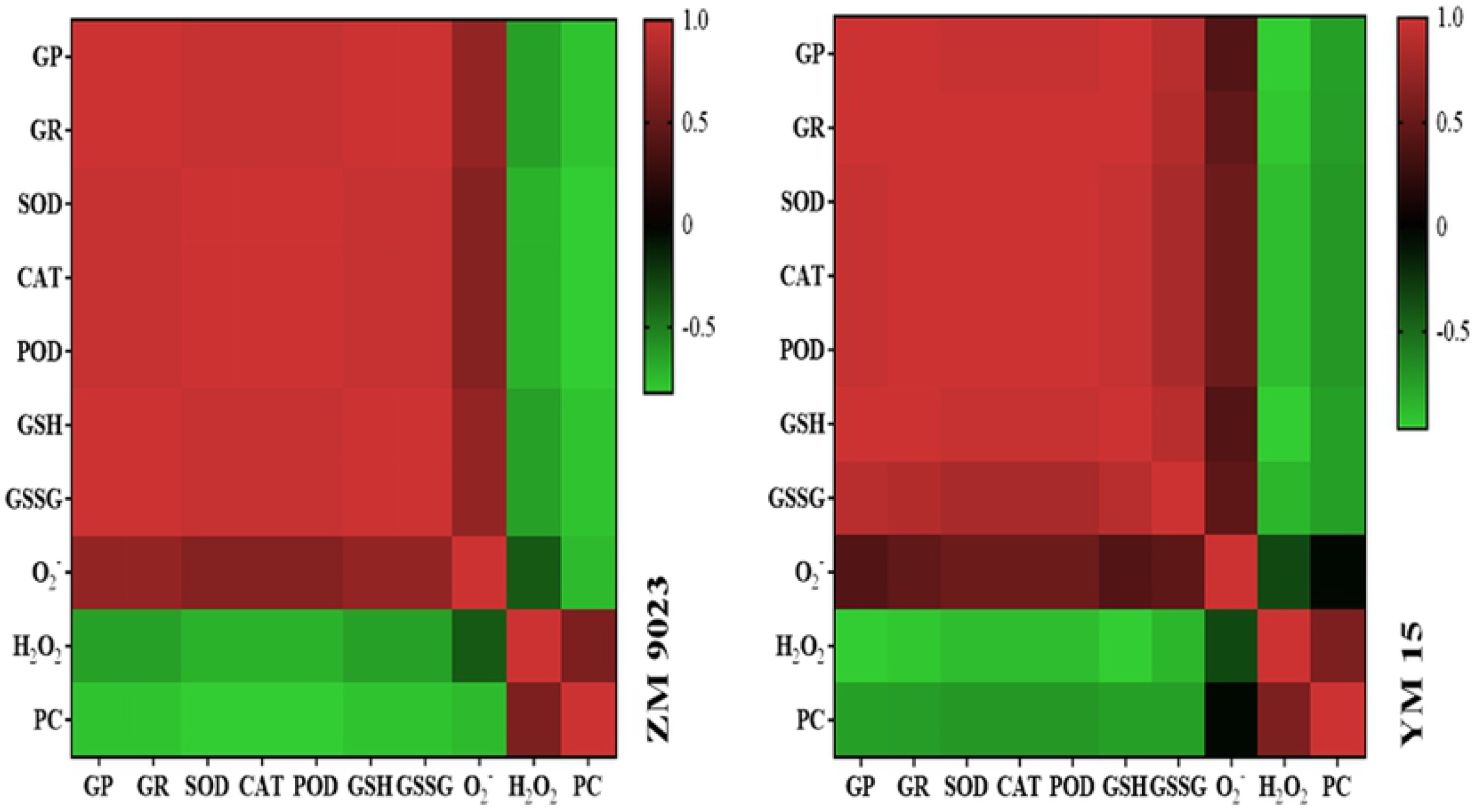
Changes in the antioxidant enzyme activities of the wheat cultivars ZM 9023 and YM 15 during artificial aging.

### Changes in non-enzymatic antioxidant contents

The content of glutathione, an important non-enzymatic antioxidant in seed, is closely related to the physiological properties of crop seeds. As shown in Fig. 5, the GSH content decreased significantly in wheat seeds of both cultivars. The GSH content in ZM 9023 immediately declined when seeds were stored under high temperature and humidity conditions, decreasing from 2,906 μg/g to 511 μg/g after 42 days. The GSH content in YM 15 decreased from 2,618 μg/g to 600 μg/g after 42 days. The same trends were observed for the GSSG contents in wheat seeds. The GSSG contents in ageing-treated seeds were significantly lower than in the control, and the artificial aging treatment decreased the GSSG contents in wheat seeds leaves by 55.26% and 75.69% compared with ZM 9023 and YM 15 controls, respectively.

**Fig.5.**
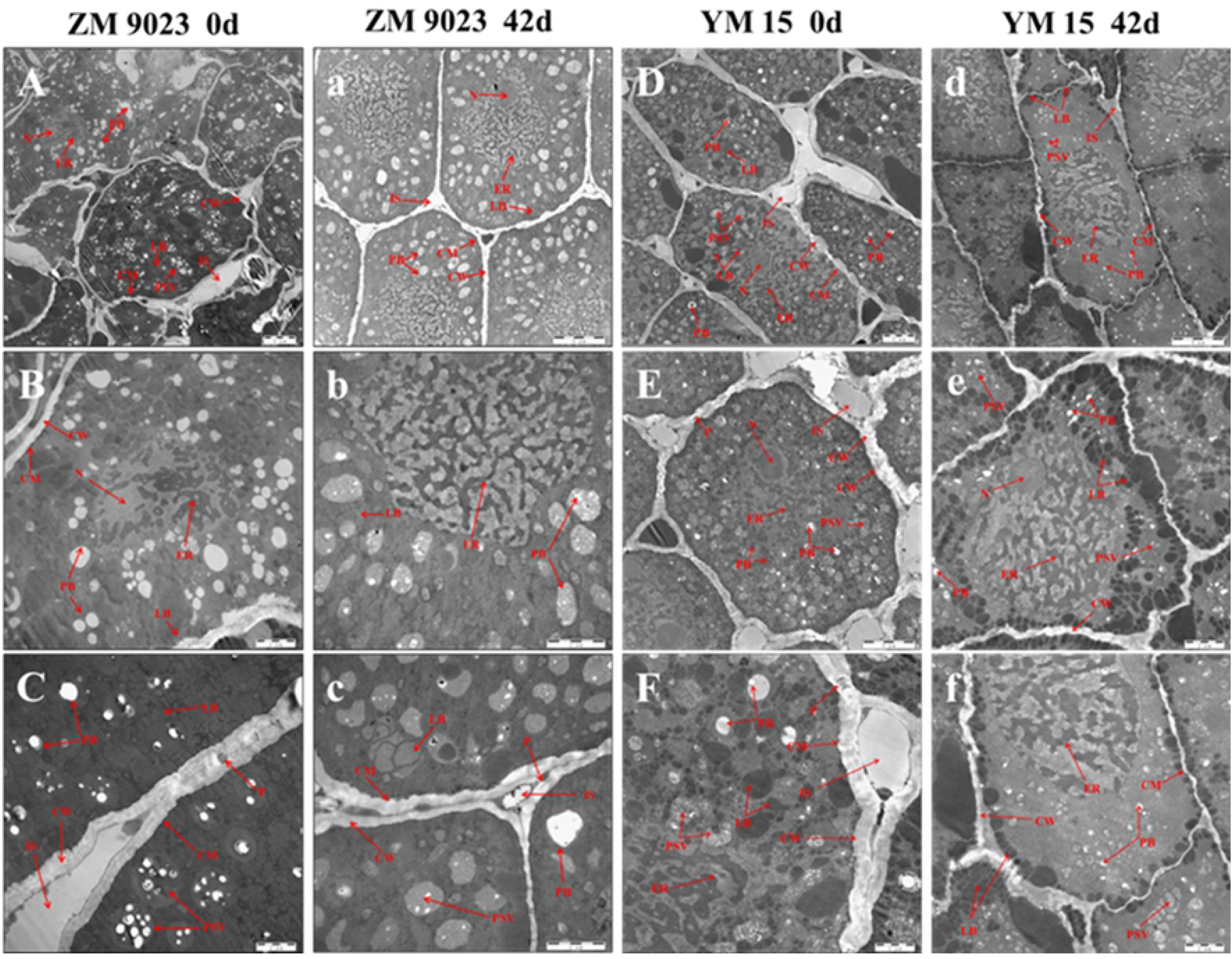
Changes in the glutathione contents of the wheat cultivars ZM 9023 and YM 15 during artificial aging.

### Correlation analyses

To further determine the relationships among seed viability, ROS content, carbonyl protein content, non-enzymatic antioxidant contents and antioxidant enzyme activities, in-depth correlation analyses were conducted using parameters of the two wheat cultivar samples that were subjected to the artificial aging treatment. The heat map shown in Fig. 6 provides an intuitive visualization of the sample information. All the germination parameters (germination rate and germination potential) were significantly and negatively correlated with the ROS and carbonyl protein contents. In addition, the carbonyl protein content was significantly and positively correlated with the ROS content. The results indicated correlations between seed vigor and carbonyl protein content.

**Fig.6.** Heat maps of the correlation between seed vigor and carbonyl protein content, reactive oxygen content, non-enzymatic antioxidant contents and antioxidant enzyme activities of wheat seeds. GR: germination rate; GP: germination potential; SOD: superoxide dismutase; CAT: catalase; POD: peroxidase; GSH: reduced glutathione; GSSG: oxidized glutathione; O_2_^−^: superoxide anion; H_2_O_2_: hydrogen peroxide; PC: protein carbonyl

### Changes in the ultrastructures of embryonic cells

The TEM analysis was conducted to observe the ultrastructures of embryos of both cultivars after 0 and 42 days of the artificial ageing treatment. As shown in Fig. 7 A–C, the ZM 9023 wheat embryo cells without the artificial ageing treatment had small and irregularly shaped nuclei. Each nucleus was surrounded by a dense, irregular endoplasmic reticulum. The cells had many protein storage vacuoles, which contained round or oval protein bodies. The protein storage vacuoles and cell membranes were surrounded by many small, dense fat bodies. The cells had a complete and smooth membrane. There were plasmodesmata and intercellular spaces between cells. However, after the artificial aging treatment, the nuclei of wheat embryo cells disappeared, the endoplasmic reticulum expanded and loosened, the protein and fat bodies in the cells were remarkably reduced, the cell membrane was damaged, and the plasmodesmata and intercellular spaces decreased significantly. In addition, the protein bodies in the cells seemed to increase in volume (Fig. 7 A–C). Similar changes were observed in YM15 wheat embryo cells. However, there was a slight difference between the two cultivars in that the protein body contents in the latter were relatively small, and the volumes of the protein bodies did not change significantly after the artificial aging treatment.

**Fig.7.** Changes in the ultrastructures of wheat embryo cells of the cultivars ZM 9023 and YM 15 during artificial aging. A–C: ZM 9023 artificial ageing treatment at 0 d; a–c: ZM 9023 artificial ageing treatment at 42 d; D– F: YM 15 artificial ageing treatment at 0 d; d–f: YM 15 artificial ageing treatment at 42 d. CM: cell membrane; CW: cell wall; ER: endoplasmic reticulum; IS: intercellular space; LB: lipid body; N: nucleus; P: plasmodesmata; PB: protein body; PSV: protein storage vacuole.

## Discussion

Wheat is a vital organism, and its vigor inevitably decreases during storage. To further explore the change mechanisms of seed vigor and their relationships with protein carbonylation during the aging process, we investigated the deterioration of vigor and the relationships among various characteristics of ZM 9023 and YM 15 during artificial aging. ZM 9023 and YM 15 are widely cultivated in China; therefore, they are representative wheat cultivar research materials.

The germination potentials and germination rates represent quality indexes that directly reflect the vigor of seeds, which are susceptible to a variety of environmental factors [5]. Simultaneously, seeds have the ability to repair themselves after exposure to external stimuli. Thus, we observed that the germination potentials and germination rates increased slightly, with fluctuations, during early storage. However, on the whole, the germination potentials and germination rates of wheat seeds significantly decreased during storage. In agreement with these results, similar negative effects of seed aging have been reported on seed vigor [2, 40].

Antioxidant enzymes and non-enzymatic antioxidants play important roles in diminishing the excess ROS produced under adverse storage conditions. They are important ROS-scavenging agents in wheat seed cells, and their decreased activity levels and contents may cause an ROS burst during the ageing process. During the artificial wheat seed aging process, the activities of SOD, CAT, POD and other antioxidant enzymes, as well as the contents of non-enzymatic antioxidants, represented by glutathione, significantly decreased, which reduced the ROS-scavenging ability and disrupted the dynamic balance between ROS production and elimination. Moreover, they also represent the first line of defense against the accumulation of carbonyl proteins by the removal of oxidatively damaged polypeptides [41].

The decline in seed vigor may result from the accumulation of ROS, which leads to the oxidative denaturation of proteins and the generation of toxic secondary metabolites that damage cellular structures during storage [42–44]. O_2_^−^ and H_2_O_2_ are common in wheat seed cells. O_2_^−^ is produced by the singlet reduction of oxygen at the end of the respiratory chain, and it reacts with Mn-SOD to generate H_2_O_2_, which may why the O_2_^−^ content did not change, whereas the H_2_O_2_ content significantly increased, during the artificial ageing process [6]. ROS may disturb mitochondrial respiration by damaging the tricarboxylic acid cycle and electron transport chain. Thus, ROS affects the energy supply of a cell’s mitochondria, resulting in the wheat seeds not having enough energy at the germination stage [45–46]. The accumulation of ROS also causes cell membrane ruptures and damages mitochondria and other organelle structures, leading to programmed cells death [47]. As observed by TEM, the cell membranes of wheat embryo cells after the artificial aging treatment were ruptured and the cells lost their selective permeability, leading to the loss of physiological functions and even death.

ROS have very strong oxidizing properties, and their accumulation in cells can cause biological macromolecules, such as proteins, lipids and nucleic acids, to be oxidized and lose physiological and biochemical functions [48]. In particular, proteins play irreplaceable roles in the cell, and oxidation is often associated with the oxidative modification of enzymes, leading to the inhibition of a wide array of enzyme activities. This may also be a reason for the decreased antioxidant enzyme activity levels in seed cells [49]. Carbonyl groups are formed on the amino-acid side chains after a protein is oxidized by ROS, and the carbonyl protein content in wheat seeds increased significantly after the artificial aging treatment [50]. The structures of proteins were changed and the original biological functions were lost, leading to cells and tissue the dysfunction. The physiological changes in the seeds eventually lead to the loss of physiological functions and even cell death, with the generation and accumulation of carbonyl proteins, which is consistent with previous studies [32, 47]

Seed storage proteins are reported to be carbonylated during storage, resulting in the seeds having lower abilities to utilize seed storage proteins for germination [21,51–52]. As observed by TEM, the protein body and lipid body contents in wheat embryo cells decreased, perhaps owing to the oxidative attack of ROS, and resulted in lipid oxidative hydrolysis reactions and protein oxidative decomposition reactions. Additionally, oxidative modifications, such as the carbonylation of proteins, lead to crosslinking between proteins, increase their volumes and change their functional properties, resulting in their inability to provide energy for seed germination. The carbonylation of mitochondrial proteins may interrupt whole-cell physiology and may be linked to progressive ageing. The carbonylation of proteins involved in organic acid metabolism, protein metabolism, energy metabolism and oxidative stress resistance may contribute greatly to mitochondrial dysfunction during storage [53–54]. Here, the germination rates and germination potentials of wheat seeds decreased during the artificial aging processing. In addition, carbonyl proteins also existed in untreated wheat seeds, perhaps owing to carbonylation occurring in wheat cells during growth, ripening, harvesting and transportation [55–56].

## Conclusion

In this study, during the artificial aging of wheat grain, the ROS production rate in wheat embryo cells was accelerated and the relative ROS-scavenging ability was weakened, resulting in ROS accumulation in the cells, which led to protein oxidation and damage to cellular structures. Protein carbonylation is a sign of protein oxidation; the higher the carbonyl protein content, the greater the degree of protein oxidation. The functional properties of proteins are altered by carbonylation, resulting in the seeds not having enough energy to sprout. Moreover, the aggregation of carbonylated proteins also leads to cell death, which ultimately leads to decreased wheat seed vigor. The results indicated that there was a negative correlation between wheat seed vigor and the carbonyl protein content. These results may provide a bases for the quality protection of wheat during storage. However, the sites of protein carbonylation during wheat seed ageing and their effects on protein functional properties still need to be further explored.

## Abbreviations

ZM 9023: Zhengmai 9023
YM 15: Yangmia 15
ROS: reactive oxygen species
O_2_^−^: superoxide anion
H_2_O_2_: hydrogen peroxide
SOD: superoxide dismutase
CAT: catalase
POD: peroxidase
GSSG: oxidized glutathione
GSH: reduced glutathione
TEM: transmission electron microscope
TTC: triphenyltetrazolium chloride

## Acknowledgements

This work was supported by the Innovative Funds Plan of Henan University of Technology (2020ZKCJ01), Cultivation Programme for Young Backbone Teachers in Henan University of Technology (21420114). We thank International Science Editing (http://www.internationalscienceediting.com) for editing this manuscript.

## Notes

The authors declare that they have no competing interests.

